# Emergence of Dual β-lactam and Colistin Resistance via bla_FRI-8_ and mcr-10.2 co-carriage on an IncFII Family Plasmid in Enterobacter vonholyi

**DOI:** 10.1101/2025.06.13.659522

**Authors:** Aneta Kovarova, Kate Ryan, Anna Tumeo, Francesca McDonagh, Christina Clarke, Martin Cormican, Georgios Miliotis

## Abstract

**Objectives:** *Enterobacter vonholyi* isolate E323169 represents a rare case of co-carriage of the antimicrobial resistance genes (ARGs) *bla*_FRI-8_, *mcr-10*.*2*. isolated from a clinical rectal swab. E323169 is one of only 12 available *E. vonholyi* genomes. To date, and across over two million pathogenic genomes scanned, only three harbour *bla*_FRI-8_ and two encode *mcr-10*.*2*, yet none exhibit both genes together. This study analyzes the genomic, phenotypic and epidemiological importance of this rare co-occurrence.

**Methods:** Species-ID for E323169 was assigned by MALDI-TOF and then re-assigned and confirmed using a multifactorial genomic workflow. Antimicrobial susceptibilities were determined by MIC assay. The genome of E323169 was sequenced on an Illumina-NextSeq-1000, assembled, and annotated for ARGs, virulence factors, and plasmid replicons detection. Comparative phylogenomics used 12 *E. vonholyi* RefSeq assemblies, and NCBI metadata were analysed for plasmid distributions of *bla*_FRI-8_ and *mcr-10*.*2*.

**Results:** E323169 carried six ARGs: four chromosomally encoded (*blaACT-91, fosA, oqxA10, oqxB9*) and two plasmid-borne (*bla*_FRI-8_ and *mcr-10*.*2*) co-located on IncFII(p14)_1_p14 replicon. Additional plasmid replicons: Col(MG828)_1 and ColRNAI_1 were also identified. By mining the NCBI Pathogen Detection pipeline, we identified *bla*_FRI-8_ on IncFII replicon in *E. asburiae* JBIWA002, and *mcr-10*.*2* on a multi-replicon (IncFIB/IncFII) plasmid in *E. kobei* 11778-yvys.

**Conclusion:** This report documents the first *E. vonholyi* isolate co-harboring *bla*_FRI-8_ and *mcr-10*.*2* on a single IncFII family plasmid, showing the widening host range of plasmid-mediated resistance to carbapenems/colistin. No prior similar co-occurrences reported, underscoring its rarity. These findings highlight the urgent need for enhanced clinical screening and genomic surveillance to curb the spread.

## 3. Main text

## 3.1. Introduction

*Enterobacter vonholyi* is a Gram-negative member of the family *Enterobacteriaceae* that does not belong to the *Enterobacter cloacae* complex (ECC). It was first isolated from retail marjoram in Germany and taxonomically classified as a novel species in 2020.^1^ As of now, only 11 *E. vonholyi* genomes sourced from human, plant, and animal isolates are available in the NCBI’s RefSeq database, with only two reports in the literature, leaving the species largely understudied. Clinical isolate E323169 represents the first detection of this species in a clinical setting in Ireland, recovered from a patient’s routine rectal swab and co-harbouring *bla*_FRI-8_ and *mcr-10*.*2* genes.

French imipenemases (FRIs) are a recently identified family of class A carbapenemases. FRI-1 through FRI-3 were first reported in European *Enterobacter* isolates between 2015 and 2017, while FRI-4 to FRI-7 were subsequently identified in Japan and Canada between 2017 and 2019.^2–6^ To date, FRI genes have been found almost exclusively in the *Enterobacter* genus, with scant evidence of horizontal transfer to other Enterobacterales. Of note, FRI genes have been identified beyond human-associated vectors, in environmental samples, suggesting cross-sectoral dissemination.^7^

The plasmid-mediated colistin-resistance (mcr) genes encode phosphoethanolamine transferases that modify lipid A and confer reduced susceptibility to colistin. Owing to their link with colistin resistance, *mcr* genes are of clinical relevance and have been widely reported, including in environmental freshwater sources in Ireland.^8^ *mcr* genes have been broadly detected in *Enterobacter* species, with the first *mcr*-10 homologue reported in 2020 on an IncFIA plasmid from a 2016 isolated clinical *Enterobacter roggenkampii* strain in China.^9,10^

The co-location of a class A carbapenemase and an *mcr* gene on the same IncFII family plasmidic contig in a clinically relevant, yet poorly characterized species, like *E. vonholyi*, which is prone to being misidentified by MALDI-TOF, could represent an epidemiological concern.^11^

### 3.2. Methods

*E. vonholyi* E323169 isolate was collected from a routine swab, initially identified, typed, sequenced and pseudo-anonymised by the National carbapenemase producing Enterobacterales (CPE) Reference Laboratory at Galway University Hospital. DNA extraction was carried using Qiagen EZ1 instrument with the EZ1/2 Tissue Kit (Qiagen, Tegelen, Netherlands) according to the manufacturer’s instructions.

Library preparation was performed using Illumina DNA Prep Kit (Illumina, USA), according to the manufacturer’s instructions, on the Biomek 4000 instrument (Beckman Coulter Life Sciences, CA, USA). Total DNA concentration was quantified with the dsDNA High Sensitivity kit and the Qubit fluorometer (ThermoFisher Scientific, Waltham, MA). 4200 TapeStation System (Agilent, CA, USA) was used to check the quality of the library prior to sequencing.

Second-generation sequencing was conducted with the NextSeq1000 system using P1 600cycle kit, (Illumina, USA).

Raw reads were filtered out using fastp v0.23.4. Shovill v1.1.0 was used for genomic assembly with setting minimum contig length to 200. Completeness and contamination were assessed using CheckM2 v1.0.2, and quality assessment was performed using QUAST v5.0.2 (Supplementary data Table S2).

Initial species identification was conducted using MALDI-ToF mass spectrometry (Bruker microflex, Bruker Daltonics GmbH, Bremen, Germany), which misidentified the isolate as belonging to ECC. Initial genomic based taxonomic delineation of E323169 was conducted using GTDB-Tk version 2.1.1 (Supplementary data Table S6). Taxonomic placement was confirmed through a multi-factorial approach combining 16S rRNA sequence analysis, digital DNA–DNA hybridization (dDDH), and Average Nucleotide Identity (ANI), with ANI values exceeding the 95% threshold relative to both the reference genome *E. vonholyi* strain STK80-C (ASM4083409v1) and the type strain E13 (ASM836455v1) (Table S8). MIC was carried out using MicroScan Gram-negative panel PNL, NEG MIC NF64 (Beckman Coulter, UK) with Mueller– Hinton II broth (Scientific Laboratory Supplies Ltd., UK). Susceptibility was determined with clinical breakpoints for Enterobacterales established by the Clinical and Laboratory Standards Institute (CLSI) M100 ED35:2025 and EUCAST breakpoints v 15.0. *E. coli* ATCC 25922 and *Pseudomonas aeruginosa* ATCC 27853 were used as controls.

All available canonical *E. vonholyi* genomes (n=11) from the NCBI RefSeq database were included for the species-wide genomic comparison (Supplementary data Table S1). Metagenome-assembled genomes (MAGs) and anomalous assemblies were excluded from the analysis. Plasmidic characteristics prediction e.g. typing, transferability, reconstruction from draft genome assemblies, was performed using MOB-typer and MOB-recon tool of MOB suite version 3.1.9.

ARGs, VFs and plasmid types were identified using Abricate v1.0.0 with CARD, VFDB and plasmidfinder databases (Supplementary data Table S3, S4 and S5). For isolate E323169, ISEScan v.1.7.3 was used to identify insertion sequence (IS) elements and DNA-features-viewer v.3.1.5 was used to visualise the genomic environment around *bla*_FRI-8_ and *mcr-10*.*2*. Core-genome based phylogenetic inference was conducted, including all RefSeq available genomes of *E. vonholyi* and using the reference genome of taxonomic neighbour *Enterobacter chengduensis* WCHECh050004 as an outgroup (Supplementary data Table S1). Roary v.3.13.0 was used to identify a core-genome of 2,925 genes. Alignment of all core-genes was conducted using Mafft v.7.407. Phylogenetic inference was conducted using RAxML v.8.2.13 with 100 bootstrap iterations, and the resulting tree was visualised with the ETE toolkit v3.1.3, with additional visualisation conducted using Scimago Graphica.

## 3.3. Results

Geographically, *E. vonholyi* genomes have been reported most frequently from Europe including Ireland, France, Germany, Switzerland and the United Kingdom. Outside Europe, three isolates originate from Assam, India, with additional detections in Bangladesh, Afghanistan, China and South Africa. Of the 11 known genomes, five derive from human hosts, and the remainder were recovered from environmental sources, plant material (including herbs) or various animal species. (Figure 1A).

**Figure 1:**
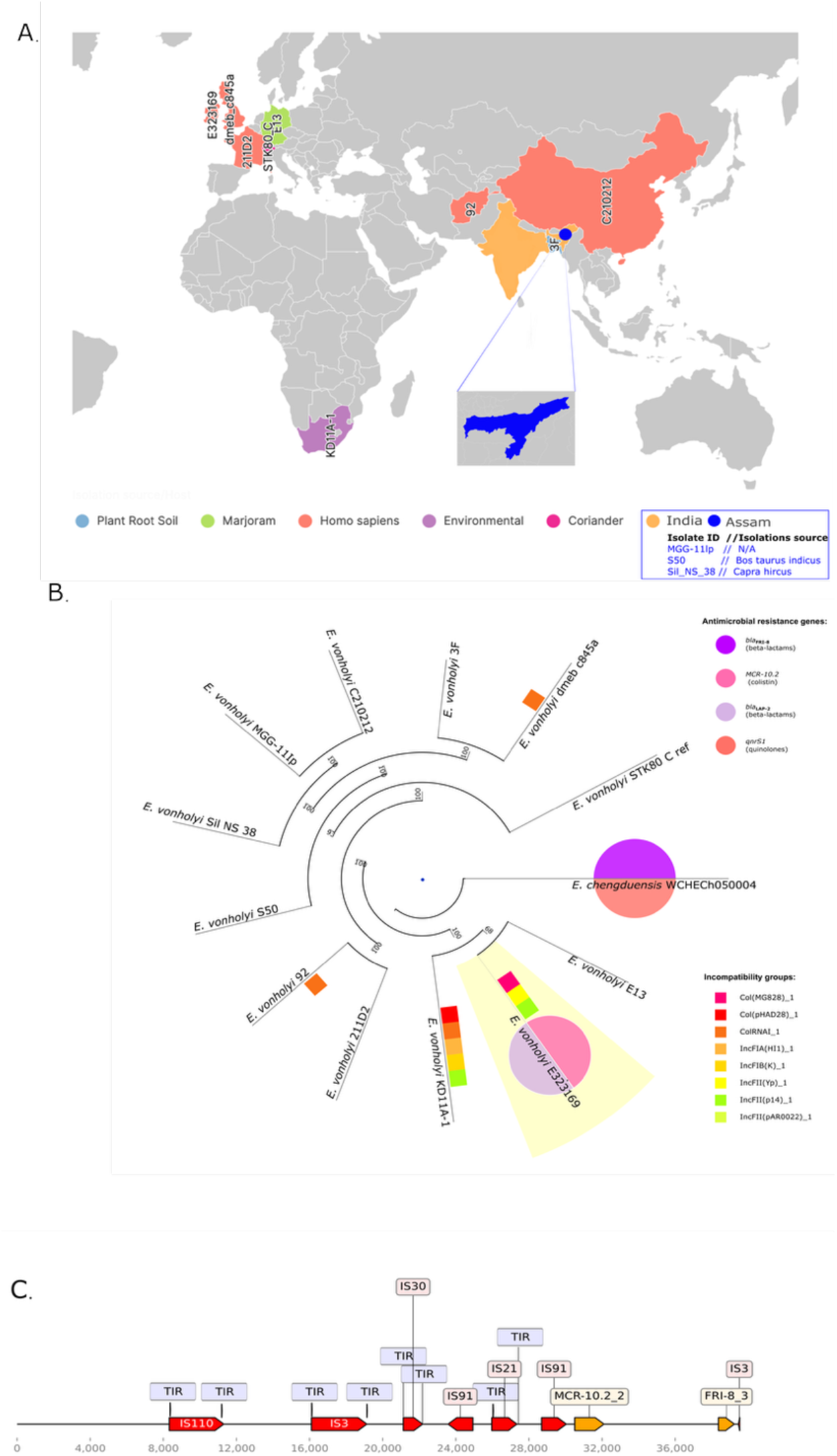
**(A)** Geo-spatial distribution of the *E. vonholyi* genomes, including colour coded - Isolation source/host and detailed description of isolates from India. **(B)** Bootstrap-supported core-genome phylogenetic analysis using outgroup comparisons. E323169 – Ireland originated isolate is highlighted in yellow, ARGs per genome are depicted in the pie charts and color coded sqaures are depicting incompatibility groups. **(C)** Linear genomic map of contig (approximately 37 000 bp) displaying ARGs (*mcr10*.*2* and *bla*_FRI_8_), IS elements and terminal inverted repeats (TIR).

Using *E. chengduensis* WCHECh050004 as an outgroup, phylogenetic reconstruction of *E. vonholyi* genomes resolved distinct lineages, reflecting a moderate level of intraspecies genetic diversity (Figure 1B). High bootstrap support (>90) underpins the close phylogenetic clustering of the *E. vonholyi* genomes S50, Sil NS 38, MCG-11lp, C20212 and 3F. In contrast, isolates S92 and umea c845a occupy separate branches, reflecting a greater genetic divergence within the main species lineage.

### 3.3.1. Minimum Inhibitory Concentration (MIC)

Minimum inhibitory concentration (MIC) testing of clinical isolate E323169 against 29 antimicrobial agents (Supplementary Table S7) demonstrated susceptibility to 24 antimicrobials. While its phenotypic response to ampicillin, cefoxitin and cefuroxime remained undetermined, E323169 was resistant to the last-resort antibiotics ertapenem and colistin. The phenotypic resistance is genetically explained by the co-carriage of *bla*_FRI-8_ and *mcr-10*.*2* genes.

### 3.3.2. Genomic analysis

#### 3.3.3.1. Isolate E323169

In total, six ARGs were identified in the genome of E323169. These include two beta-lactamase genes, namely the AmpC-type beta-lactamase *bla*_ACT-91_, and the class A carbapenemase *bla*_FRI-8_. The isolate also carried *fosA*, which is a fosfomycin resistance encoding gene, and *mcr-10*.*2*. Additionally, the presence of *oqxA10* and *oqxB9* genes was noted, suggesting a potential for reduced susceptibility to quinolones. The resistance genes *bla*_FRI-8_ and *mcr-10*.*2* were identified on the same plasmid-borne contig, while the remaining four ARGs (*bla*_ACT-91_, *fosA, oqxA10*, and *oqxB9*) were located on chromosomal contigs.

Only three VFs, *csgG, entB*, and *ompA*, were detected in this isolate, all located on chromosomally derived contigs. Three plasmid replicons were also identified in isolate E323169: Col(MG828)_1, ColRNAI_1, and IncFII(p14)_1_p14. ColRNAI_1 was detected on chromosomal contigs, likely reflecting algorithmic limitations in distinguishing plasmid sequences within draft genome assemblies.

#### 3.3.3.2. Database mining and species wide comparative analysis

Using AMRFinderPlus to screen all >2 million bacterial assemblies in NCBI’s Pathogen Detection pipeline (April 2025), we detected *bla*_FRI-8_ in three *ECC* genomes, and in none of the non-ECC genomes: clinical *E. cloacae* 2022GO-00182 (USA), clinical *Enterobacter asburiae* N21-01785 (Canada), and environmental *E. asburiae* JBIWA002 (lake water, Japan). Similarly, only two Chinese clinical *Enterobacter kobei* isolates (11778 and 11778-yvys) were found to harbor *mcr-10*.*2*, emphasizing the scarcity of both these genes among pathogenic taxa.

In isolate E323169, both *bla*_FRI-8_ and *mcr*-10.2 are predicted to reside on the same IncFII(p14)_1_p14 plasmid replicon. Examination of plasmid’s contig00014 (Figure 1C) shows each gene flanked by distinct, incomplete insertion sequences lacking terminal inverted repeat, suggesting potential for independent mobilization. Specifically, *bla*_FRI-8_ is adjacent to a truncated IS3 element (38 360–39 256 bp), while *mcr*-10.2 is bordered by a partial IS91 sequence (30 499– 32 118 bp).

Analysis of the 11 *E. vonholyi* genomes in RefSeq revealed a highly conserved ARG repertoire (Figure 2A,B). Aside from the plasmid-borne resistance determinants in clinical isolate E323169, all other detected ARGs were chromosomally encoded: every strain carried *fosA*, and *mcr*-10.1 was found on the chromosome of isolate KD11A-1.

**Figure 2:**
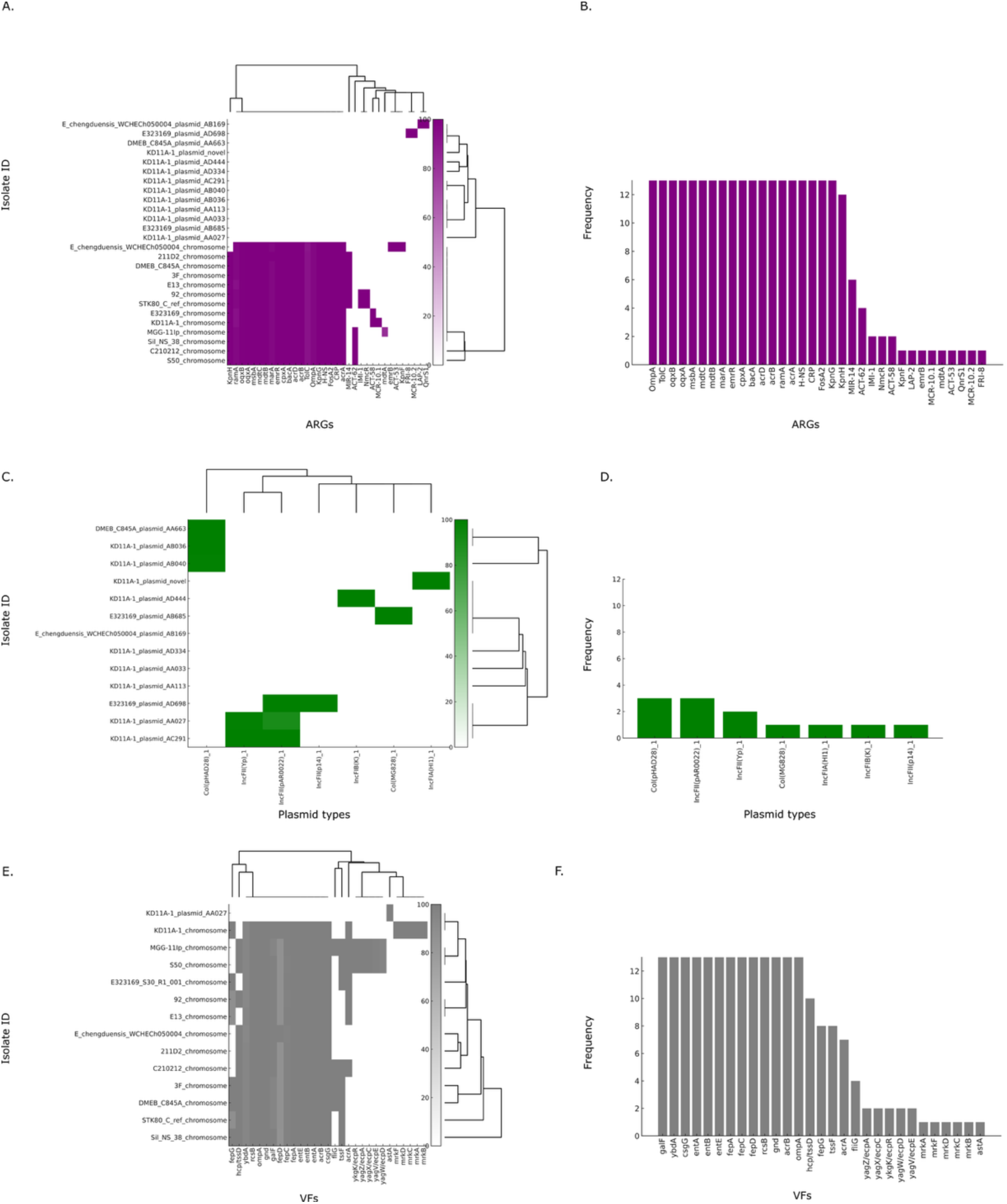
**(A)** Distribution of ARGs identified in *E. vonholyi* genomes and the outgroup isolate *E. chengduensis*, classified by chromosomal or plasmid localization. **(B)** The frequency of each ARG present across all isolates in this study. (C) Presence of plasmid types across all isolates in this study. (D) The frequency of each plasmid type present across all isolates in this study. (E) Distribution of VFs identified in *E. vonholyi* genomes and the outgroup isolate *E. chengduensis*, classified by chromosomal or plasmid localization. (F) The frequency of each VF present across all isolates in this study.

Plasmid typing analysis identified seven distinct plasmid replicon types across the *E. vonholyi* dataset (Figure 2C, D), with replicons detected only in the genomes of E323169, KD11A-1 and DMEB_C845A.

Overall, VF profiles were largely conserved across isolates, with a few exceptions. The *mrk* operon, encoding type 3 fimbriae, was detected only in the environmental isolate KD11A-1, whereas the *yag* operon, responsible for ECP pilus biogenesis, was found exclusively in isolates MGG-11Ip and S50, originating from an unidentified environmental source and an animal host, respectively (Figure 2E,F). Notably, the only acquired VF across the species was the EAST-1 enterotoxin gene (*astA*), located on a plasmid in KD11A-1.

## 3.4. Discussion

This genomic analysis of *E. vonholyi* present a geographically widespread, genetically diverse species with emerging multidrug-resistance and variable genomic virulence potential.

Its cross-sectoral recovery from both clinical and environmental/animal sources across Europe and parts of Asia and Africa suggest a capacity for broad host–niche colonization. Phylogenetically we identified a core clade alongside divergent branches, suggestive of both recent clonal spread and deeper intraspecies diversification.

The clinical isolate E323169 typifies the species’ capacity to acquire high-impact resistance determinants. Distinct IS elements flanking each ARG point to independent mobilization events, mirroring reports of *bla*_FRI-8_ in other *Enterobacter* species and *mcr-10* in *E. kobei*.^12^ Among the three ECC *bla*_FRI-8_–positive isolates, the fully resolved genome of *E. asburiae* JBIWA002 carries *bla*_FRI-8_ on an IncFII plasmid replicon. Of the two *mcr-10*.*2*–harbouring strains, the complete genome of isolate 11778-yvys encodes *mcr-10*.*2* on a multireplicon plasmid bearing both IncFIB and IncFII backbones.^13^ Such multireplicon plasmids, common in *Enterobacteriaceae* and strongly associated with ARGs, feature broad host-range adaptability, specialized stability modules, and robust maintenance systems that facilitate long-term persistence in bacterial populations.^14^

By contrast, VFs remain largely chromosomal and conserved; the *mrk* operon (type 3 fimbriae) and *yag* operon (ECP pilus) are each restricted to specific environmental sublineages, and only isolate KD11A-1 carries the plasmid-encoded heat-stable enterotoxin EAST-1 encoding gene *astA* associated with diarrheagenic *E. coli*.^15^

Overall, *E. vonholyi* combines a largely conserved core genome with sporadic uptake of mobile resistance and enterotoxin elements. The co-emergence of IncFII family plasmid-borne *bla*_FRI-8_ and *mcr-10*.*2* is suggestive of the need for continued genomic surveillance to better understand this non-ECC *Enterobacter* species.

## 2. Abbreviations

NCBI: National Center for Biotechnology Information
mcr: plasmid-mediated colistin-resistance
FRI: French imipenemases
CPE: Carbapenemase producing enterobacterales
MALDI-ToF: Matrix-Assisted Laser Desorption/Ionisation Time-of-Flight
ANI: Average nucleotide identity
MIC: Minimal Inhibition Concentration
ARGs: Antimicrobial resistance genes
VFs: Virulence Factors
MAGs: Metagenome-assembled genomes
ECC: *Enterobacter cloacae* complex
dDDH: DNA-DNA hybridization
CLSI: Clinical and Laboratory Standards Institute
TIR: Terminal inverted repeats
IS: Insertion sequences

## 4. Data availability

Complete genome sequence of *Enterobacter vonholyi* E323169 have been deposited in the GenBank database; under BioSample accession number SAMN47937951, genome accession JBNCAP000000000 and BioProject number PRJNA1250442.

## 5. Conflicts of interest

The authors declare that there are no conflicts of interest.

## 6. Ethics approval

No ethics approval required.

## 7. Funding

Funding statement: The research conducted in this publication was funded by 1) Taighde Éireann – Research Ireland under grant number GOIPG/2023/4515; 2) the EPA Research Programme 2021–2030 as Government of Ireland initiative funded by the Department of the Environment, Climate and Communications; 3) Science Foundation Ireland (SFI) under Grant number 18/CRT/214.

## 8. Acknowledgements

Thanks to University Hospital Galway Microbiology Department for facilitating access to MALDI-TOF for species identification, and to the National CPE Reference Laboratory at Galway University Hospital for providing the sample, identifying, typing, sequencing and pseudo-anonymising the isolate.

## 9. Limitations

*E. vonholyi* can be easily misidentified as a member of the *Enterobacter cloacae* complex (ECC) on MALDI-TOF MS due to the close genetic and protein profile similarities within the complex, which often lead to ambiguous or incorrect species-level identifications.

## Notes

### Competing Interest Statement

The authors have declared no competing interest.

